# Electrophysiological characterization of human atria: the understated role of temperature

**DOI:** 10.1101/2020.12.07.414573

**Authors:** Rupamanjari Majumder, Afnan Nabizath Mohamed Nazer, Alexander V. Panfilov, Eberhard Bodenschatz, Yong Wang

**Affiliations:** Laboratory for Fluid Physics, Pattern Formation and Biocomplexity, Max Planck Institute for Dynamics and Self-Organization, 37077 Göttingen, Germany; German Center for Cardiovascular Research, Partner Site Göttingen, 37075 Göttingen, Germany; University Medical Center Göttingen, 37075 Göttingen, Germany; Arrhythmia Department, Almazov National Medical Research Centre, Saint Petersburg, Russia; Department of Physics and Astronomy, Ghent University, Krijgslaan 281, S9 Gent 9000, Belgium; Laboratory of Computational Biology and Medicine, Ural Federal University, Ekaterinburg, Russia; Laboratory of Atomic and Solid-State Physics and Sibley School of Mechanical and Aerospace Engineering, Cornell University, Ithaca, New York 14853, USA

**Keywords:** cardiac electrophysiology, temperature, remodelling, human atria, atrial fibrillation, action potential, restitution

## Abstract

Ambient temperature has a profound influence on cellular electrophysiology through direct control over the gating mechanisms of different ion channels. In the heart, low temperature is known to favour prolongation of the action potential. However, not much is known about the influence of temperature on other important characterisation parameters such as the resting membrane potential (RMP), excitability, morphology and characteristics of the action potential (AP), restitution properties, conduction velocity (CV) of signal propagation, etc. Here we present the first, detailed, systematic *in silico* study of the electrophysiological characterization of cardiomyocytes from different regions of the normal human atria, based on the effects of ambient temperature (5 −50°*C*). We observe that RMP decreases with increasing temperature. At ∼ 48°*C*, the cells lose their excitability. Our studies show that different parts of the atria react differently to the same changes in temperature. In tissue simulations a drop in temperature correlated positively with a decrease in CV, but the decrease was region-dependent, as expected. In this article we show how this heterogeneous response can provide an explanation for the development of a proarrhythmic substrate during mild hypothermia. We use the above concept to propose a treatment strategy for atrial fibrillation that involves severe hypothermia in specific regions of the heart for a duration of only ∼ 200ms.

## 1 INTRODUCTION

Abnormal generation and propagation of electrical signals in the heart lead to disturbances in its regular electromechanical pump function. Such disturbances can occur abruptly over a few beats and disappear by themselves, or they can persist for a sufficiently long period of time to be classified as an arrhythmia. Cardiac arrhythmias can be of different types. The most lethal variants include ventricular fibrillation (VF), ventricular tachycardia (VT) and atrial fibrillation (AF), of which persistent AF has the highest rate of occurrence in human patients (Nattel, 2002). Unlike VT and VF, AF is treatable by pharmacological methods or by other means, such as radiofrequency (RF) ablation (Hussein et al., 2016). The method involves destruction of excitable heart tissue to *(i)* remove the “source” of the arrhythmia, and *(ii)* to prevent recurrence at the same site. Despite its high success rate, RF ablation involves a major compromise, namely the infliction of permanent damage to healthy heart tissue. Therefore, alternative, low-energy defibrillation methods, are in great demand. In particular, recently, optogenetics has emerged as a promising tool to study and control wave dynamics in cardiac tissue (O’Shea et al., 2019; Nyns et al., 2019; Nussinovitch and Gepstein, 2015; Entcheva, 2013; Bingen et al., 2014; Bruegmann et al., 2010). However, despite its success in the treatment of arrhythmias in small mammalian hearts (Bruegmann et al., 2010; Bingen et al., 2014; Nyns et al., 2019; Majumder et al., 2018), optogenetics suffers from two major limitations: (*i*) the long term effects of genetic modification are not completely known, which makes clinical translation debatable; and (*ii*) penetration of light in cardiac tissue is very poor, making it difficult to defibrillate deep tissue (Bruegmann et al., 2016).

Ambient temperature (*T*) has long been known to influence cellular and molecular processes (Hannon, 1958; Bjørnstad et al., 1993; Hodgkin and Katz, 1949; Kiyosue et al., 1993; Zhao and Boulant, 2005; Kågstroöm et al., 2012). In particular, temperature has a direct bearing on ion channel kinetics, through its influence on the rate constants of gating, and reaction rates of certain sub-cellular chemical processes, which thereby allow cells to produce action potentials with altered morphology, characteristics and cell-to-cell propagation speeds. In cardiac electrophysiology, the effect of temperature is modelled by dividing the time constant for opening/closing of an ion channel, by a temperature-dependent scaling factor *Q*(*T*), (or equivalently, multiplying the rate constants of the relevant sub-cellular chemical reaction, by *Q*(*T*)). *Q*(*T*) has a power-law dependence on temperature, according to the Arrhenius law of heat transfer (Hodgkin and Katz, 1949; Malki and Zlochiver, 2018). Previous studies showed that in small mammals such as the rabbit and ground squirrel, lowering temperature leads to increase in the resting membrane potential (RMP), reduction in action potential amplitude (APA), and prolongation of action potential duration (APD) at 50, 70, and 90% repolarization, at the single cell level (Fedorov et al., 2005), whereas, in tissue, signal conduction velocity (CV) was found to decrease with decrease in temperature (Fedorov et al., 2008). These features resemble those of optogenetically modified cells, with temperature replacing light and eliminating the need for a genetic modification.

In a recent study, Tobón et al. (2013) presented a model for anatomically realistic human atria, based on geometry and fiber data acquired using Diffusion Tensor Magnetic Resonance Imaging (DTMRI). Their model characterizes the atria while taking into account, the differences in action potential morphology and characteristics, and conduction anisotropy, as measured from different regions of the atria. We developed a model of the electrophysiologically-detailed human atria, based on the Courtemanche-Ramirez-Nattel (CRN) model for single cell action potentials (AP) from human atrial cardiomyocytes (Courtemanche et al., 1998), using the AP-characteristic data that was used to develop the Tobón model (Tobón et al., 2013). We incorporated temperature dependence into the kinetics of the different ion channels in the different atrial regions, and used it to demonstrate low-temperature induced break-up of scroll waves during mild hypothermia in accordance with the results of Filippi et al. (2014) for canine hearts and the low-temperature induced suppression of scroll waves during severe hypothermia, a finding that leads us to consider thermal control as a possible tool for arrhythmia management in the future.

## 2 METHODS

We modeled the time evolution of the transmembrane potential *V* according to Eq. 1, where we described the net ionic current (*I*_*ion*_) flowing across the cell membrane, according to the CRN model (Courtemanche et al., 1998) for human atrial cardiomyocytes.

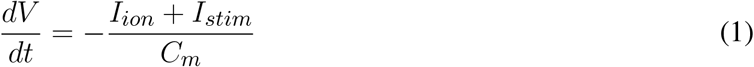

Here, *C*_*m*_ is the specific capacitance of the cell membrane (in *µ*F*/*cm^2^). As given in Eq. 2, *I*_*ion*_ represents the sum of 12 ionic currents: the fast *Na*^+^ current (*I*_*Na*_), the inward rectifier *K* ^+^ current (*I*_*K*1_), the transient outward *K*^+^ current (*I*_*to*_), the ultra-rapid *K*^+^ current (*I*_*Kur*_), the rapid and slow delayed rectifier *K*^+^ currents (*I*_*Kr*_ and *I*_*Ks*_, respectively), the *Na*^+^ and *Ca*^2+^ background currents (*I*_*BNa*_, and *I*_*CaL*_), the *Na*^+^*/K*^+^ pump current, the *Ca*^2+^ pump current (*I*_*pCa*_), the *Na*^+^*/Ca*^2+^ exchanger current (*I*_*NaCa*_) and the *L* − *type Ca*^2+^ current (*I*_*CaL*_).

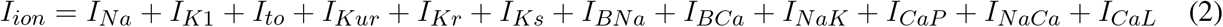

We used forward Euler method to solve Eq. 1 with a time step of 0.02ms. In order to model the action potential during chronic AF remodelling, a combination of parameters sets from earlier publications (Courtemanche et al., 1999; Pandit et al., 2005; Zhang et al., 2005) was adopted. In particular, we down-regulated the maximal conductances of *I*_*to*_, and *I*_*CaL*_ by 85%, and 74%, respectively, increased the maximal conductance of *I*_*K*1_ by 250%, increased the time constant for activation of *I*_*CaL*_ by 62%, shifted the fast *Na*^+^ inactivation curve by +1.6mV, the *I*_*to*_ activation curve by +16mV, and the *I*_*CaL*_ activation curve by −5.4mV.

We took into account the following regions of the atria (check Fig. 2 for reference): the crista terminalis (CT), Bachmann’s bundle (BB), pectinate muscles in the right atrial free wall (PM), the fossa ovalis (FO), isthmus of the right atrium (RIS), the atrial appendages (APG), the atrio-ventricular regions (AVR), sinoatrial node (SAN), pulmonary veins (PV) and finally, the atrial working myocardium (AWM). To model the region-based heterogeneity of the atria, we adjusted the maximal conductances of specific ion channels. The adjustment was constrained by the condition that in switching from normal to remodelled atria, the APD_90_ values remained within ∼ 10% deviation of the ones obtained by Tobón et al. (2013). The full list of maximal conductances is given in Table 1. Fig. 1 shows a comparison between APs obtained from different regions of the atria, as produced using our model in both control and remodelled atria.

**Table 1.** Region-specific adjustments to the CRN model, with corresponding APD90 and CV at 37 °*C*, in the control atria.

|  | CT | BBFO | PM | PV | RIS | AWM | AVR | APG |
| --- | --- | --- | --- | --- | --- | --- | --- | --- |
| $G_{K1}$ (nS/pF) | 0.09 | 0.117 | 0.1188 | 0.117 | 0.117 | 0.117 | 0.117 | 0.09 |
| $G_{to}$ (nS/pF) | 0.1652 | 0.1652 | 0.1652 | 0.1652 | 0.1652 | 0.1652 | 0.1652 | 0.0578 |
| $G_{Kr}$ (nS/pF) | 0.0294 | 0.0294 | 0.0588 | 0.0294 | 0.0294 | 0.0294 | 0.1176 | 0.0324 |
| $G_{CaL}$ (nS/pF) | 0.1238 | 0.1238 | 0.1238 | 0.1238 | 0.1238 | 0.1238 | 0.1176 | 0.0619 |
| $APD_{90}$ (ms) | 307 | 284 | 242 | 284 | 284 | 284 | 183 | 253 |
| $\mathcal{D}$ ($cm^2/ms$) | 0.009 | 0.0064 | 0.0064 | 0.0016 | 0.0013 | 0.0025 | 0.0025 | 0.0025 |
| CV (cm/s) | 142.5 | 117.75 | 117.75 | 54.75 | 48 | 69.75 | 69.75 | 71.25 |

**Figure 1.**
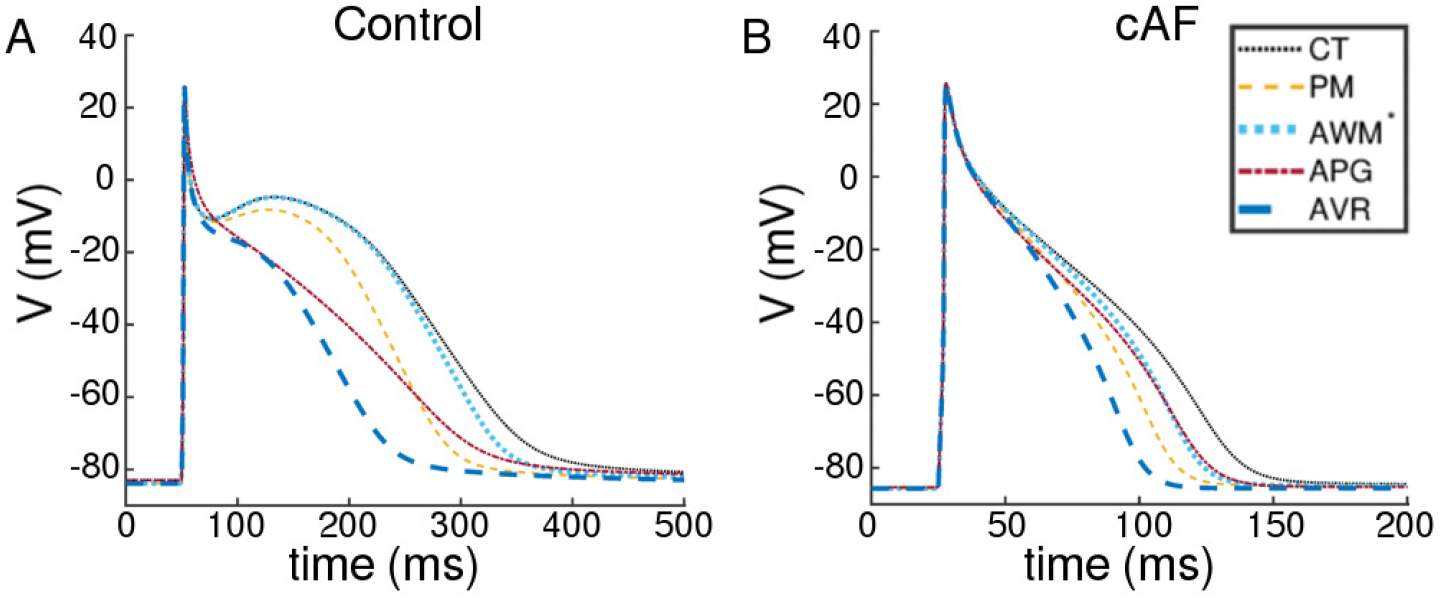
Action potentials obtained using region-specific models for different parts of the atria, under (A)control and (B) chronic AF (cAF) remodelled conditions. The dotted black line represents the AP fromcrista terminalis (CT), the dashed orange line represents the AP from the pectinate muscle region (PM); thedashed maroon line represents the AP from left and right atrial appendages (APG), and the thick dashedblue line represents the AP from the left and right atrioventicular regions (AVR). The rest of the atria, i.e,the Bachmann’s bundle (BB), fossa ovalis (FO), right and left atria, pulmonary veins (PV) and isthmus ofthe right atrium (RIS), is modelled using a single set of parameters. The corresponding AP is shown usinga thick dotted blue line (AWM *).

**Figure 2.**
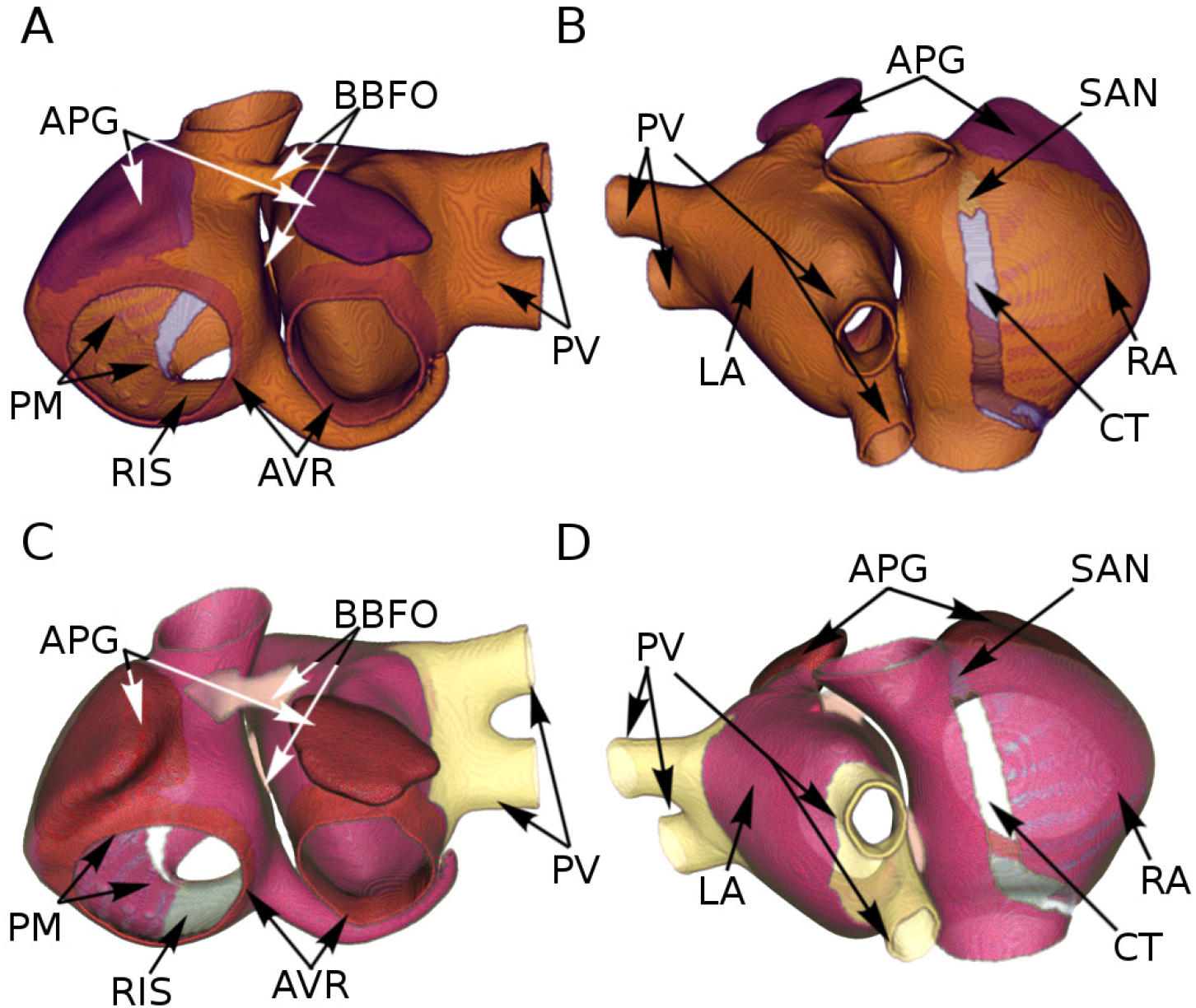
Segmentation of the atria according to AP-based (upper panel) and conduction-based (lower panel) heterogeneity.

Like Tobón et al. (2013), we considered differences in propagation velocities in different parts of the atria. Thus, the idea was to get regions with propagation speed as fast as 120 −140cm*/*s (PM and CT) coexist with regions of propagation, as slow as 25 −45cm*/*s (SAN and RIS). To this purpose, we extended our cell model to two dimensions (2D), whereby the regional CV could be factored in through the choice of the diffusion coefficient 𝒟. As in case of the region-specific values of the ion channel conductances, 𝒟, was determined based on the constraint that in switching from normal to remodelled atria, region-specific CV demonstrated acceptable resemblance to the ones reported by Tobón et al. (2013), in both normal and remodelled cases. In 2D, spatiotemporal evolution of *V* was modelled according to the following reaction-diffusion-type equation:

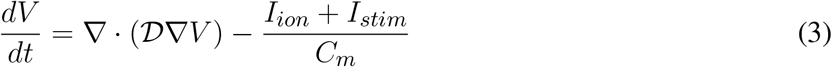

The temporal part was solved numerically using forward Euler method, as in single cells, whereas the Laplacian in the spatial part (𝒟 is a constant in 2D) was solved using centered finite difference technique, with a spatial resolution of 0.03cm, which was fixed by the spatial resolution of the DTMRI data, used for the anatomically realistic simulations.

Next, we incorporated temperature scaling of parameters in this model, according to the Hodgkin and Katz formalism Hodgkin and Katz (1949), by either multiplying the rate constants *α*(37°*C*) and *β*(37°*C*) of the gating variables of specific ion channels (see below) by *Q*(*T*), or, by dividing the time constant *τ* (37°*C*) of these ion channels by *Q*(*T*) (in case the *α* and *β* values were unavailable).

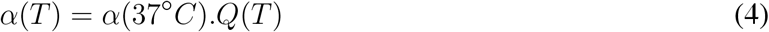

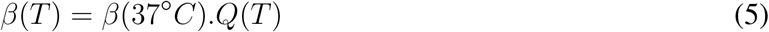

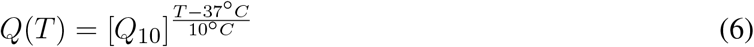

Th *Q*_10_ factors used for the different currents, were obtained from Malki *et al*. Malki and Zlochiver (2018) *(see Table 2 below)*.

**Table 2.** Temperature scaling factors (*Q*_10_ values) for different ionic currents

| Current | $I_{Na}$ | $I_{to}$ | $I_{Kur}$ | $I_{Kr}$ | $I_{Ks}$ | $I_{CaL}$ |
| --- | --- | --- | --- | --- | --- | --- |
| $Q_{10}$ | 3.0 | 2.2 | 2.2 | 3.3 | 2.57 | 2.2 |

We studied RMP in single cells. To measure its temperature dependence, we started with the standard steady-state conditions of the CRN base model. We altered the maximal conductances of specific ion channels, as described in Table 1 and set ambient temperature to *T*_1_ (5°*C < T*_1_ *<* 50°*C*). We then allowed the system to evolve in time for 10s, in the absence of an applied stimulus. The minimum current (*χ*), required to elicit an AP at different temperatures, was measured in a manner, similar to RMP measurements, with the stimulus being applied at the end of 10s, when the system was assumed to have reached a more-or-less steady state.

To measure restitution characteristics, we used a 2D stripe domain containing 512 × 10 grid points. After allowing the system to stabilize for 5s at a given ambient temperature *T*_1_, we initiated a train of 10 right-propagating plane waves (paced at 1.0Hz), from the region *x <* 5 grid points, by applying current stimuli of strength 25*µ*A and duration 2.0ms. After the application of the 10^*th*^ stimulus, we let the system evolve freely for a brief period of time (the diastolic interval DI), and applied a final (11^*th*^) pulse of the same strength and duration. We repeated the procedure at 5°*C* ≤ *T*_1_ ≤ 50°*C* in steps of 5°*C*, for all considered regions of the atria. In each case, we measured APD_50_, APD_90_ and propagation speed (CV), from the wave of electrical activation, corresponding to the 11^*th*^ stimulus.

Lastly, we studied the effect of change in temperature on scroll wave dynamics in the anatomically-realistic, electrophysiologically-detailed human atria. To construct the geometry and fiber structure, we used open-source DTMRI data available from Tobón et al. (2013). The irregular boundaries were handled by a phase field implementation (Fenton et al., 2005; Clayton and Panfilov, 2008), whereby, we embedded the atria in a rectangular parallelepiped grid structure composed of regular cubic nodes. We defined a field variable *ϕ* in this domain, such that *t* = 0, all points outside the atria are reset (*ϕ* (*x, y, z*) = 0) and all points belonging to the atria, are set (*ϕ* (*x, y, z*) = 1). Next, we evolved *ϕ* in space and time using a relaxation method (Eq. 7) to smoothen the interface between the atrial boundary and the exterior.

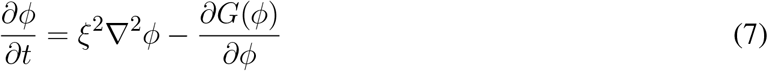

The parameter *ξ* determines the width of the interface, and *G*(*ϕ*) is a standard double-well potential:

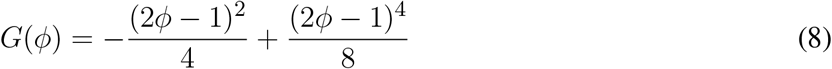

We used the spatial profile of *ϕ*, obtained after 500 iterations (with a spatial resolution of 0.03cm and time step *δt* = 0.02ms), for studying electrical activity in the atria. At each node with *ϕ* ≠ 0, we integrated the temperature sensitive ionic models of the region-specific cardiomyocytes, that we developed within the first part of the paper. For the pacemaker, however, we used the AWM model with reduced CV, and a simple periodic step function to represent the nodal electrical activity from the designated region. Then, using an 18 - point stencil, we solved Eq. 9, a modified version of Eq. 3, to compute the spatiotemporal evolution of *V*.

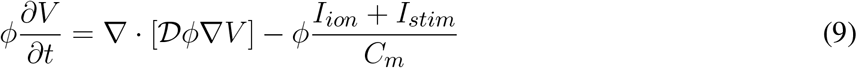

Note, that in these 3D anatomical models, the diffusion coefficient 𝒟 is no longer a scalar constant, but a rank-3 tensor, whose components (*d*_*ij*_) are related to the fiber directions (*a*_*i*_), as follows:

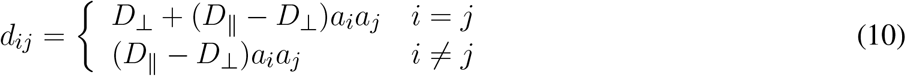

The atria were segmented according to AP-based heterogeneity as per Fig. 2A-B. To incorporate conduction heteroheneity, the atria were segmented according to Fig. 2C-D.

To compute the temperature field (*T*) in 2D and 3D simulations, we solved the time-dependent heat-transfer equation (Paul et al., 2014):

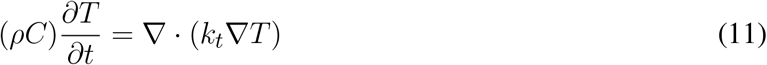

Here, *ρ, C*, and *k*_*t*_ represent, respectively, the density (1050kg*/*m^3^), specific heat capacity (4219J*/*(kgK)), and thermal conductivity (0.7W*/*(mK)) of human cardiac tissue. For these simulations, we assumed that the temperature both outside the atria, and in the cavities, was 37°*C*.

## 3 RESULTS

### 3.1 Single cell simulations

Our simulations of electrical activity in cardiomyocytes isolated from different parts of the atria, show that RMP correlates negatively with ambient temperature for both normal (Control) and remodelled (cAF)

conditions. This is shown in Fig. 3A-B. As a direct consequence, at elevated temperatures (*>* 48°*C*), cardiomyocytes from all regions of the atria become functionally inexcitable. Fig. 3C-D illustrate the temperature dependence of the minimum amount of electric charge (*χ*, measured in Coulombs) required by the cell to produce an action potential in both Control and cAF cases. At *T* ≳ 48°*C*, irrespective of the value of *χ*, one cannot obtain action potentials because the RMP is extremely low, leading to complete inactivation of the fast *Na*^+^ channels, which are responsible for the primary excitability of the cell. The applied excitation then dissipates exponentially, as in a decaying RC-circuit. Exposure to such high temperatures for up to several seconds can even lead to permanent damage of the cell membrane, with complete loss of excitability (Kolandaivelu et al., 2010).

**Figure 3.**
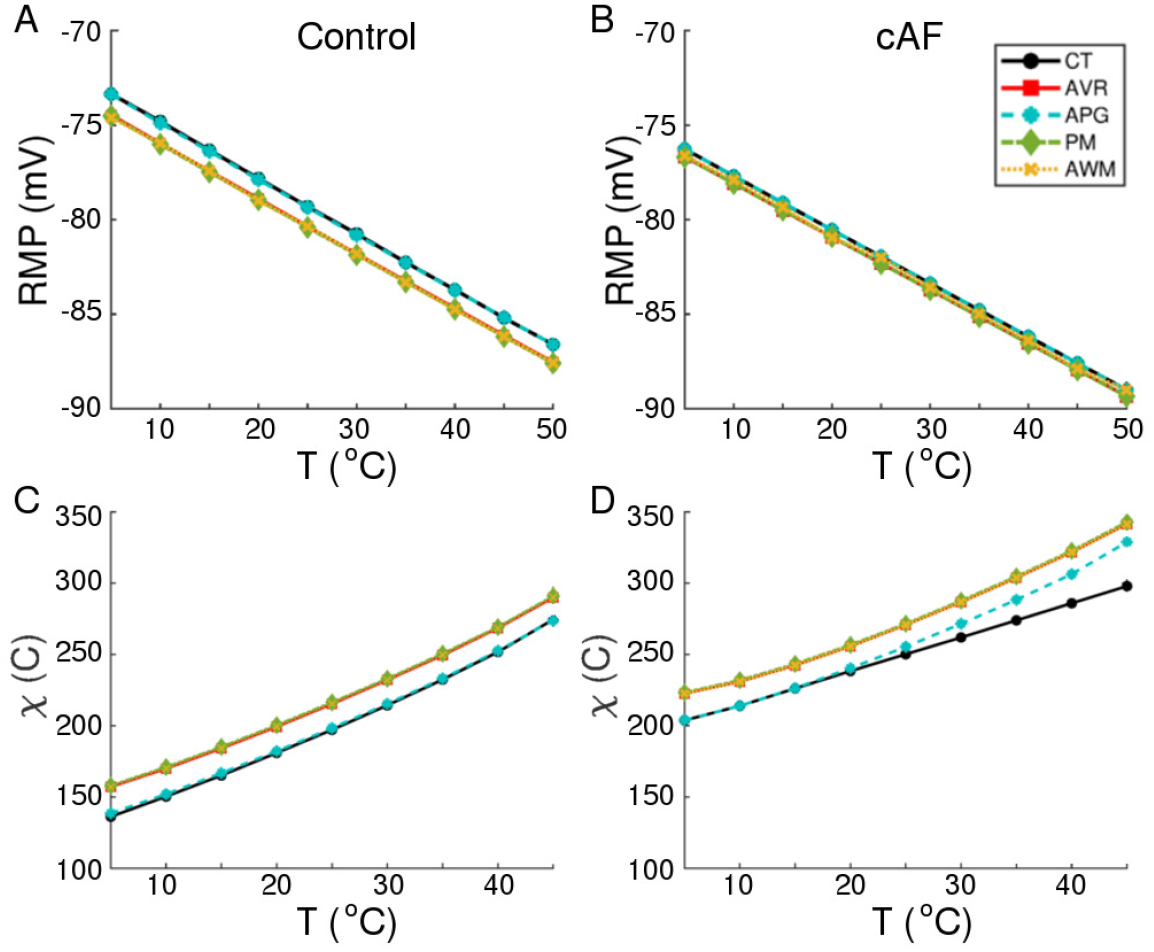
Temperature-dependence of RMP, and minimum stimulus (*χ*, in Coulombs) required to produce an action potential, in normal (A, C) and remodelled (B, D) cardiac cells, isolated from different regions of the human atria (CT, AVR, APG, PM and AWM *).

Our model illustrates (in consonance with what is known about human atrial cardiomyocytes), that at body temperature (37°*C*), the action potential is longest (∼ 307ms) in the CT region, with a pronounced notch around ∼ −10mV. The AP is shortest in the AVR region (∼183ms), with a triangular morphology and complete disappearance of the notch. AP morphology is also triangular in the AWM region, whereas, in APG and PM, a short plateau is visible. AP morphology exhibits dramatic dependence on temperature. Comparative AP traces from different parts of the atria at different temperatures (5° −50°*C*) reveal that, the qualitative effect on AP morphology is most pronounced in the CT region, where lowering of temperature is accompanied by a prolongation of the plateau phase. The least dependence on temperature is observed in the APG and AVR regions, where the AP is generally less affected, except within the temperature range 20°*C* ≲ *T* ≲ 30°*C*, where alternans appears. For quantitative comparison, temperature-dependency curves for APD_90_, APD_50_ and notch depth (*d*_*notch*_) are presented in Fig. 4 A-D. Clearly, the effect of temperature on APD_90_ is maximum in the CT region (see Fig. 4A), followed by BBFO, AWM, PV and RIS (comparable with PV). In APG and AVR, while a decrease in *T* from 50°*C* shows an initial increase in APD_90_ similar to other atrial regions, below a critical temperature (∼ 25°*C* for AVR and ∼ 15°*C* for APG), APD_90_ decreases with decrease in *T*.

**Figure 4.**
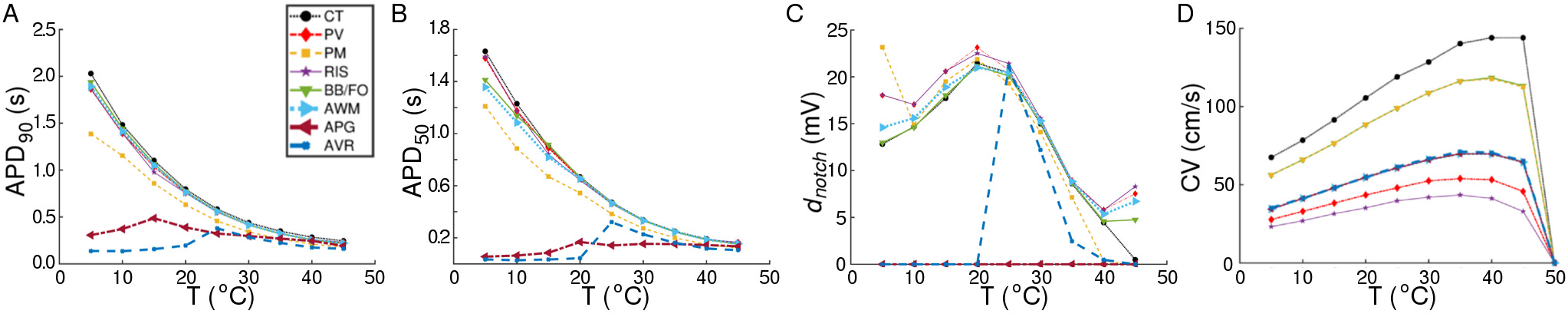
Comparison of region-specific temperature-dependencies of AP characteristics (A) APD_90_, (B) APD_50_, and (C) notch depth (*d*_*notch*_), and (D) conduction velocity (CV) under normal (Control) conditions.

We observed that at these critical temperatures, the single-cell models for APG and AVR showed alternans. Similar trends were observed with APD_50_ (see Fig. 4B, although, in this case the APG model remained approximately independent of temperature at *T* ≳20°*C*. Below ∼ 20°*C*, APD_50_ decreased with decrease in *T*. Finally, in Fig. 4C, we illustrate the temperature-dependence of *d*_*notch*_. Our results demonstrate that for all regions except APG and AVR, the AP exhibits a notch at body temperature (37°*C*). The notch depth grows as *T* is decreased down to ∼20°*C*. Below this temperature, *d*_*notch*_ decreases monotonically except at PM, PV and RIS, where, it goes through a minimum at ∼ 10°*C*. When heated to temperatures above 37°*C, d*_*notch*_ decreases everywhere except at APG and AVR. Here, heating can be associated with a gradual triangulation of the AP morphology. In all regions except CT, *d*_*notch*_ goes through a second minimum at ≃ 40°*C*, followed by a slow increase with further rise in temperature. The AP from APG is always triangular, i.e, *d*_*notch*_ = 0mV at all temperatures, whereas the notch disappears from AVR APs at *T* ≲ 20°*C*. At this temperature, *d*_*notch,AV R*_ ≈ *d*_*notch,RIS*_.

### 3.2 Conduction velocity studies

A study of temperature-dependence of conduction velocity (CV), as measured from different parts of the atria, showed that CV increased linearly with increase in *T* for 5°*C* ≲*T* ≲ 25°*C*. Above 25°*C*, CV still increased with T, but at a rate that became progressively slower. Finally, above 40°*C*, CV decreased rapidly to 0 by 50°*C*, at which point, action potentials were no longer formed. This is shown in Fig. 4 D. Normalization of CV by its value at 37°*C* yielded a family of curves, each with slope ∼1.02. This is interesting, given that the models used to simulate the different regions of the atria, make use of different electrophysiological parameters. The temperature-dependence of APD_90_ and CV in single cardiac cells and stripe domains constructed from cells belonging to different regions of the atria are tabulated in Table 3.

**Table 3.** Comparison between region-specific APD_50_, APD_90_ and CV values for control (C) and remodelled (R) atria, at 37°*C*, from tissue simulations.

|  | Condition | CT | BBFO | PM | PV | RIS | AWM | AVR | APG |
| --- | --- | --- | --- | --- | --- | --- | --- | --- | --- |
| APD<sub>50</sub> (ms) | C | 223.74 | 218.58 | 184.02 | 220.02 | 218.82 | 221 | 134.24 | 154.64 |
| APD<sub>50</sub> (ms) | R | 73.3 | 66 | 60.4 | 62.8 | 61.4 | 65.3 | 52.76 | 60.3 |
| APD<sub>90</sub> (ms) | C | 319.48 | 294.4 | 253.62 | 296.52 | 296.38 | 296.44 | 213.98 | 276.1 |
| APD<sub>90</sub> (ms) | R | 107.4 | 92.5 | 85.7 | 90.6 | 90 | 91.7 | 77.1 | 94.2 |
| CV (cm/s) | C | 142.5 | 117.75 | 117.75 | 54.75 | 48 | 69.75 | 69.75 | 71.25 |
| CV (cm/s) | R | 137.25 | 113.6 | 113.25 | 51.75 | 40.9 | 67.5 | 67.5 | 68.6 |

### 3.3 APD and CV restitution

In our studies with single atrial cells, we noted that electrical properties of the cell at any given temperature, depended heavily on the frequency with which the cell was stimulated. In particular, for regions such as CT, BBFO, AWM, PV and RIS, which show APs ∼ 𝒪 (2s) at 5°*C*, it is important to understand how the sub-cellular processes cope with slow and fast periodic electrical stimulation. Therefore, we investigated the temperature-dependence of restitution. Our results are presented in Fig. 5 (A-H). We observed that, although the APD restitution curve for CT at 37°*C* follows the standard trend, with maximum slope *<* 1.0, the restitution behaviour is affected remarkably at high and low temperatures. In particular, at *T <* 37°*C*, the APD restitution curve shows an initial slow rising phase (with very small positive slope). Then, at a critical diastolic interval (*DI*_*c*_), the slope changes dramatically, as if the restitution behaviour is undergoing some sort of state transition. Finally, at large DI, APD saturates to a large value. Fig. 6A-C show detailed analysis of the APD restitution curve for CT at different temperatures.

**Figure 5.**
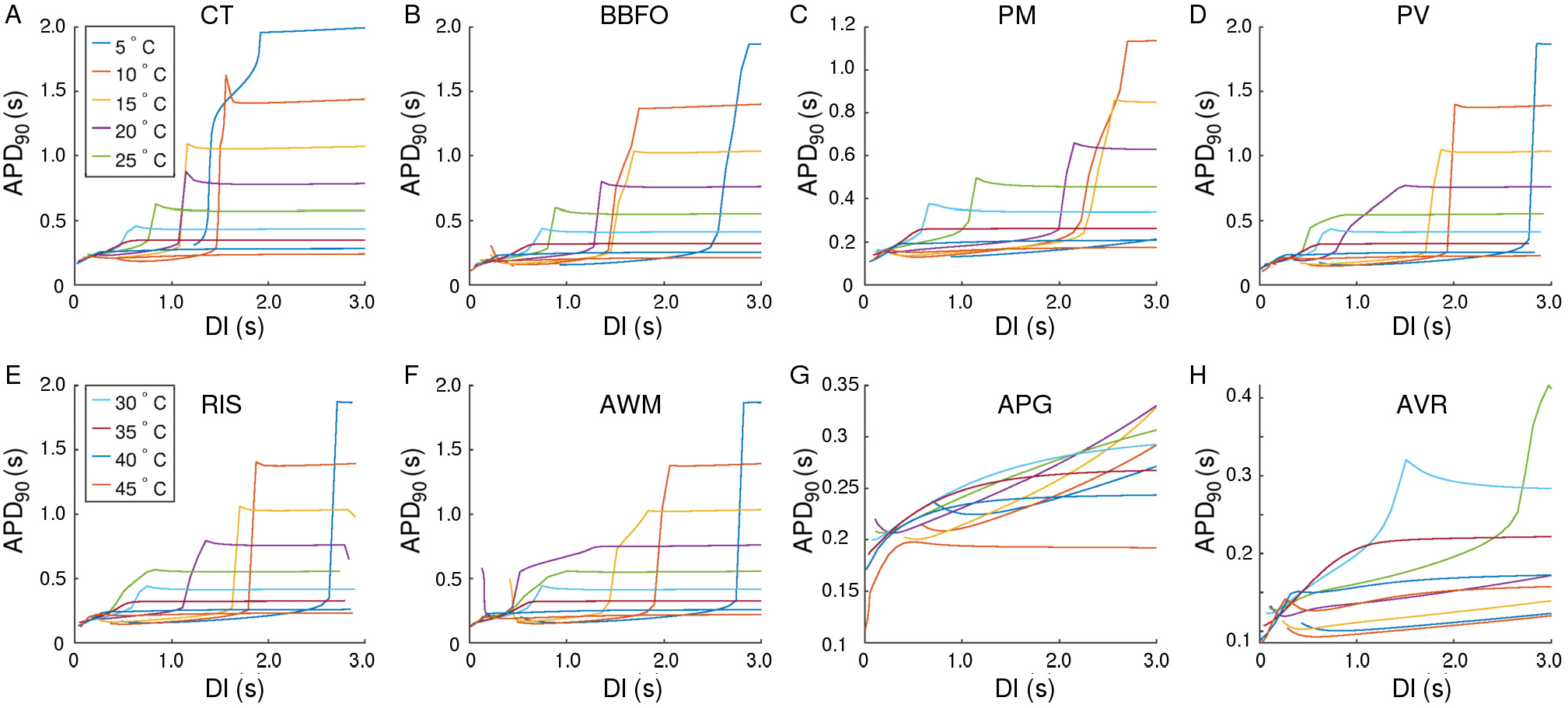
Temperature-dependence of APD_90_ restitution curves recorded from (15.36 mm ×3 mm) tissue stripes containing cells from CT, BBFO, PM, PV, RIS, AWM, APG and AVR, respectively. Restitution was measured using the dynamic S1-S2 protocol, whereby, each tissue stripe was first allowed to reach a steady state at a given temperature (5° *< T <* 45°*C*). Then a series of 20 S1 pulses were applied at the left end of the stripe, at a pacing frequency of 1 Hz. Finally, a 21^*st*^ stimulus (S2) was applied in a similar manner, but after a variable diastolic interval (DI). Our results indicate the occurrence of a phenomenon similar to a first order state transition, at very low temperatures in most atrial regions. At high temperatures (*T >* 40°*C*), the APD restitution curves exhibit a negative slope.

**Figure 6.**
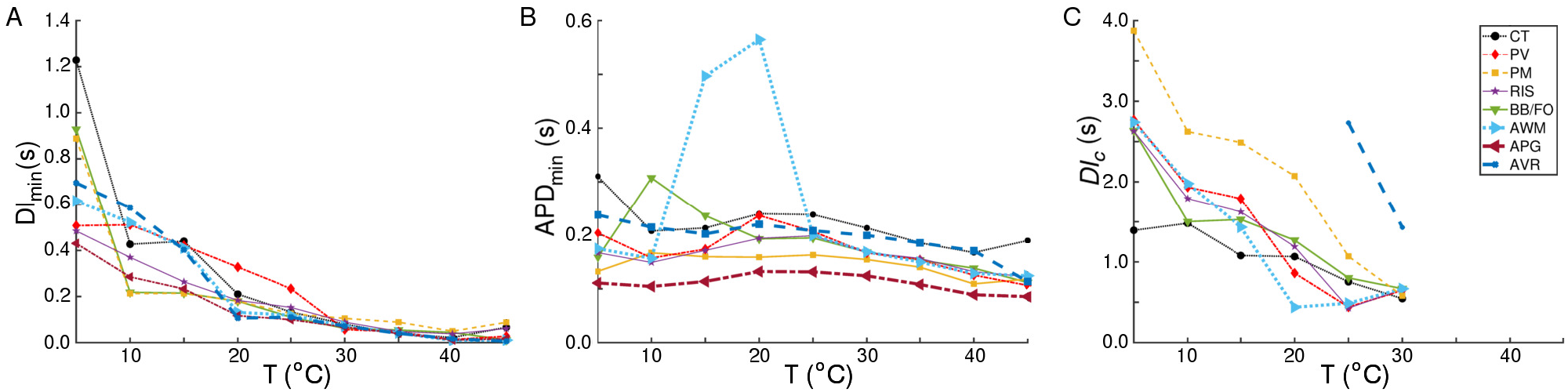
Temperature-dependence of (A) DI_*min*_, (B) APD_*min*_, and (C) *DI*_*c*_, as measured from CT, BBFO, PM, PV, RIS, AWM, APG and AVR. For all 3 parameters, we observe similar trends in the temperature dependency curves. The large deviation in the trend of APD_*min*_ around 15° −20°*C* for AWM shows up because of the atypical behaviour of the APD restitution curves, measured at these temperatures (see Fig. 5F). Similarly, the temperature dependency curve for AVR in (C) shows deviations from the general trend of *DI*_*c*_ in all the other regions. This is because the APD restitution curves for AVR do not show signatures of the state transition at other temperatures. Furthermore, the APD restitution curves for APG do not undergo this transition at any temperature. Thus, the *DI*_*c*_ − *T* curve for APG is absent in (C).

The APD resitution curve for AWM shows similar trends as CT, except that at temperatures ∼15° −20°*C* the CT-like curves are preceded by an additional rapid phase, in which the restitution curve has a large negative slope. The detailed analysis of the APD restitution curve for AWM is presented in Fig. 6A-C. APD restitution trends similar to CT were also observed in PM, PV, BBFO and RIS regions, although, as presented in the analysis plots of Fig. 6, the maximum value of the APD reached by a system, and the steepness of the slope during the possible state transition, increased with decrease in temperature. Finally, the APD restitution curves of APG and AVR exhibited behaviours that were different from the rest of the atria. In APG, at *T <* 35°*C*, APD saturation did not occur, even at DI= 3000ms. The curves then showed an initial phase with a small negative slope, which gradually increased to a positive value. Hereafter, APD kept increasing with DI. At *T* ≳ 35°*C*, the APD restitution curves followed the standard trend with saturation of APD at large DI. As for AVR, the APD restitution curves exhibited trends similar to APG at all temperatures, except ∼25° −30°*C*. In this range of temperatures, the APD restitution curves showed signatures of a possible state transition.

In most atrial regions, the CV restitution curves exhibited a temperature-dependency. However, CV, itself, appeared to be independent of DI in all regions except CT, PM and BBFO. These data are presented in the plots of Fig. 7.

**Figure 7.**
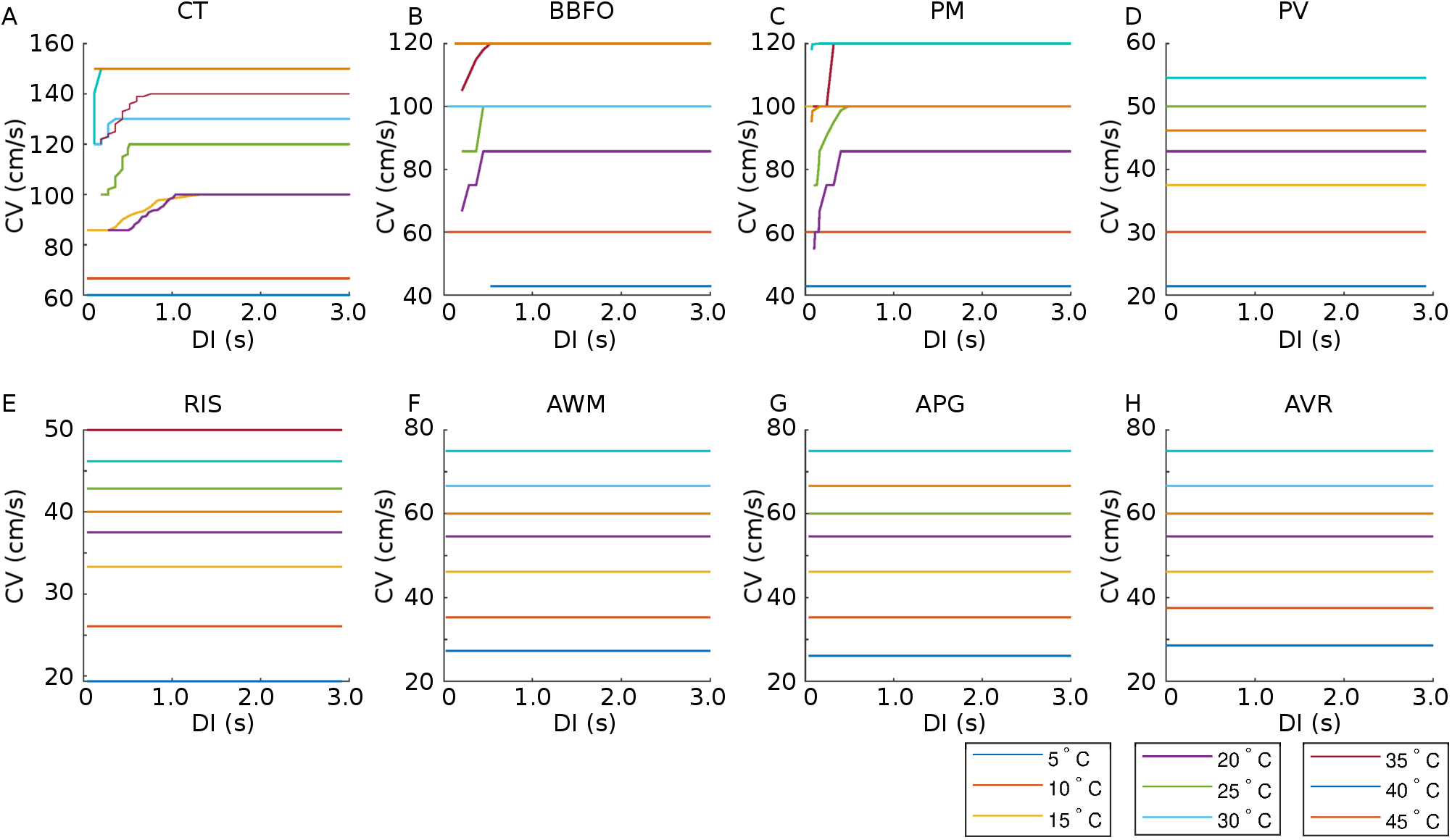
Temperature-dependence of CV restitution curves recorded from (15.36 mm × 3 mm) tissue stripes containing cells from CT, BBFO, PM, PV, RIS, AWM, APG and AVR, respectively. Restitution was measured using the dynamic S1-S2 protocol (see Methods for details). Our results show that CV is more or less independent of DI for most regions (PV, RIS, AWM, APG and AVR), at all temperatures. However, in some regions (CT, BBFO and PM), CV exhibits initial fluctuations at low DI at moderate temperatures.

### 3.4 Effect of cooling the anatomically realistic human atria

Based on our findings at the single cell level and with stripes, we investigated the effect of cooling on anatomically realistic, electrophysiologically detailed human atria, in the presence of scroll waves. First, we simulated wave propagation in the 3D atrial model at normal body temperature (37°*C*). We looked at the propagation patterns of electrical signals originating at the SAN, and compared our findings with those of Tobón et al. (2013). As illustrated in Fig. 8, the electrical signal initiated at the SAN propagated down the right atrium with a triangular wavefront, due to large differences in conduction velocities between the CT and AWM regions. In our model of the normal heart, the time to full depolarization of both atria was ∼140ms, which is in line with the observations of Tobón et al. (2013). In the case of cAF, due to the reduction in signal wavelength, repolarization of the right atrium sets in before the left atrium is fully depolarized. These results are presented in Fig. 8.

**Figure 8.**
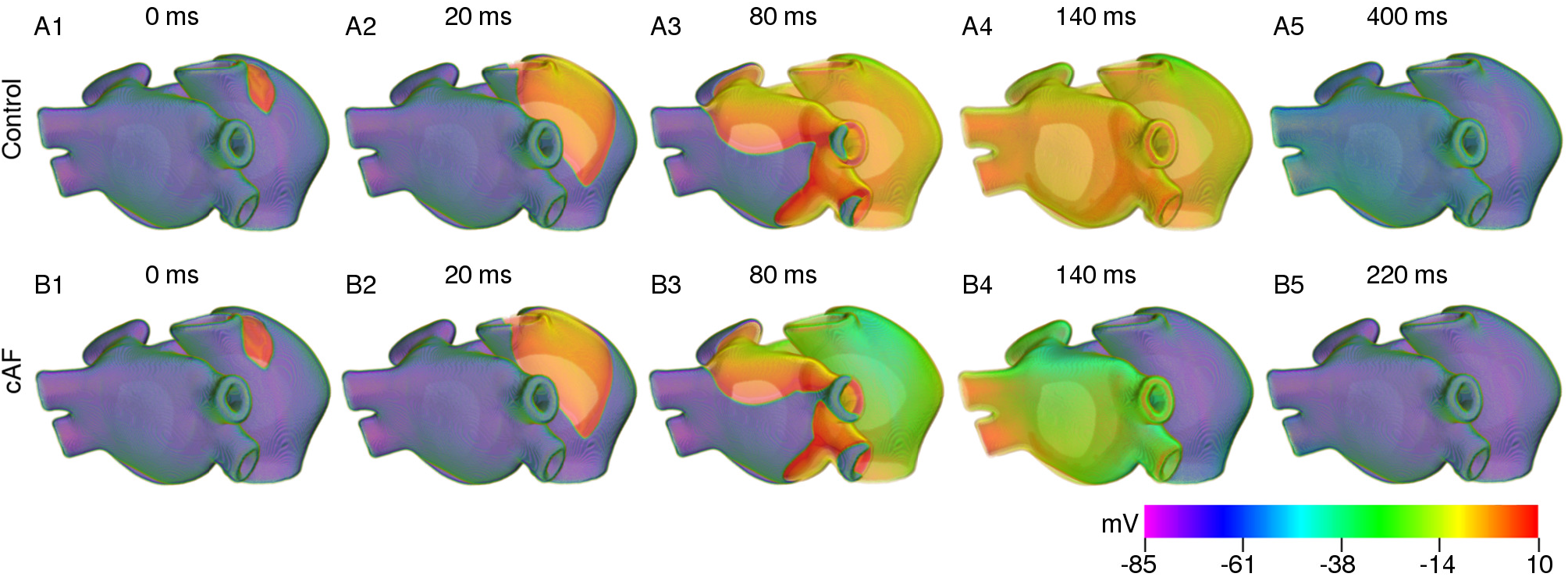
Propagation of a plane wave through the human atria at normal body temperature (37°C)

We produced a pair of counter-rotating scroll waves in the remodelled human atria (see Fig. 9) by applying S1-S2 cross-field protocol. The S1 wave originated at the SAN. When it had propagated sufficiently into the atria, such that the tissue behind the wave tail had already started to recover from refractoriness, we applied the S2 stimulus, perpendicular to the S1 wave. This resulted in the production of a pair of counter-rotating scroll waves, of which, one (centered in the right APG region) became the principal driver of the rotational activity, whereas the other anchored to one of the caval veins and converted itself into anatomical reentry.

**Figure 9.**
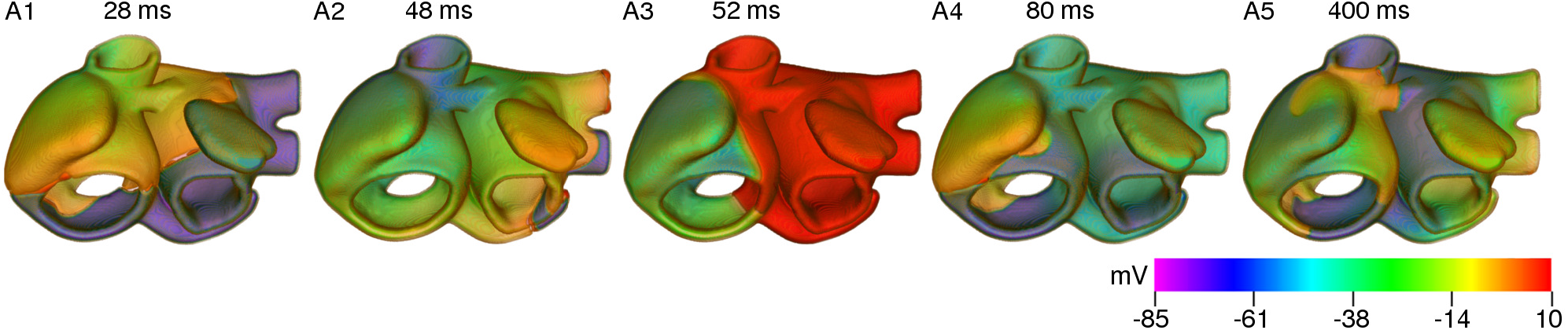
Formation of a pair of counter-rotating scroll waves by S1-S2 cross-field protocol at 37°*C*.

In order to test the effect of cooling on the spatiotemporal dynamics of scroll waves in the atria, as a first approach, we applied uniform cooling to the entire thickness of the atrial wall, in all parts of the atria. As proven by our simulations with the stripe domain, different atrial regions responded differently to the same decrease in ambient temperature. A direct consequence of this heterogeneous electrophysiological response, is the increased heterogeneity of AP characteristics, which happens with reduction in temperature. At moderately low temperatures (20°*C < T <* 35°*C*), the arrhythmia stabilized via establishment of a stable core of the rotating scroll wave (see Fig. 10 A). At lower temperatures (10°*C < T* ≲20°*C*), alternans in some parts of the atria promoted degeneration of a single scroll wave into multiple self-sustained wavelets that coordinate to maintain a state of sustained AF (see Fig. 10 B & C). Interestingly, further decrease in temperature to ∼5°*C* led to suppression of the scroll waves, due to substantially reduced excitability of the tissue, which could no longer sustain the scroll wave (see Fig. 10 D). In this situation, the drop in temperature caused the filament of the primary scroll wave to meander away from the right APG region into the right AVR, where it got absorbed at the boundary. This further led to termination of the reentrant wave around the caval vein, which received its driving force from the scroll wave in the right APG. Thus severe cooling for very short duration, within 200 ms as shown in our study, can form the basis for a new approach to arrhythmia management and therapy.

**Figure 10.**
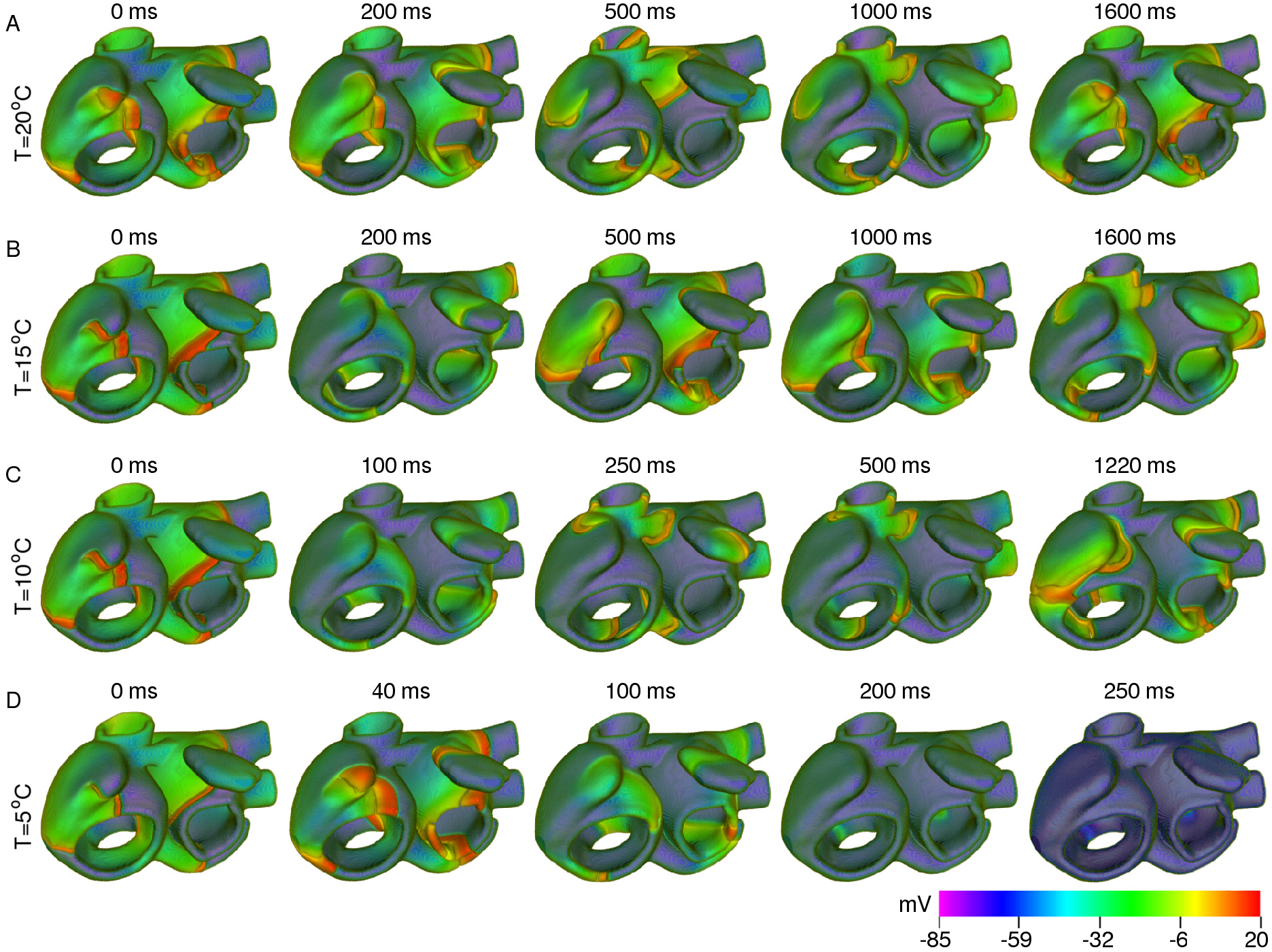
Scroll wave dynamics in the remodelled human atria, under the effect of global cooling at temperatures (A) 20°*C*, (B) 15°*C*, (C) 10°*C* and (D) 5°*C*. Moderate cooling (15 −20°*C*) leads to stabilization of the initial scroll wave, with reduced period. At ∼10°*C*, the scroll wave breaks up. The atria then supports multiple scroll waves that maintain a state of sustained AF. Severe cooling (∼ 5°*C*) leads to suppression of the scroll wave, within 200ms.

## 4 DISCUSSION

We performed an extensive, systematic thermal characterization of the anatomically-realistic, electrophysiologically-detailed human atria. Our study gives novel insights into the crucial role of temperature in the determination of spatiotemporal dynamics of scroll waves in the atria, when subjected to cooling at different temperatures. By constructing stripe domains that contain cardiomyocytes specific to different atrial regions (e.g, CT, PM, PV, BBFO, RIS, AWM, APG and AVR), we demonstrated how lowering of temperature from 37°*C* (normal body temperature) to ≃20°*C* (room temperature) could already induce alternans in different regions of the atria, thereby facilitating the development of a proarrhythmic substrate. Our results indicate that further cooling (≃10°*C*) could lead to break up of stable rotating scroll waves into multiple wavelets that were capable of sustaining themselves through mutual interaction. Finally, severe cooling (∼ 5°*C*) for very short duration (∼200ms) could successfully terminate all electrical activity in the atria, through destabilization caused by sudden, substantial decrease in tissue excitability.

Very little is known about the effects of temperature on spiral and scroll wave dynamics in cardiac tissue, and even less so, in human subjects. In our study at the single cell and tissue level, we observed increase in the RMP with decrease in temperature. Similar increase was observed by Fedorov et al. (2005) in hearts of rabbits and ground squirrels, who also reported a 10-fold decrease in CV, and increased excitation threshold in going from 37°*C* −3°*C*. In contrast, our studies revealed that in human atria, the decrease in CV (ΔCV) could reach up to a maximum of ∼ 3 −*fold* in the CT region for a decrease in temperature from 37°*C* −5°*C*. ΔCV was found to be less in other regions of the atria, for the same decrease in temperature. We also observed an increased excitation threshold, with decrease in temperature, consistent with the findings of Fedorov et al. (2005).

In another experimental study using hearts of rabbits and ground squirrels, Egorov et al. (2012) reported observing spatially discordant alternans at temperatures as low as 17°*C*. Similar alternans was also observed in canine ventricular preparations by Fenton et al. (2013), who provide a quantitative description of emergent proarrhythmic properties of restitution, conduction velocity, and alternans regimes as a function of temperature. We observed APD alternans at low temperatures (15°*C* −20°*C*) in specific regions of the atria, which led to the promotion of wavebreaks in the simulations involving anatomically realistic atria. Furthermore, we observed similar temperature-dependency trends of the APD restitution curves as Fenton et al. (2013) at lower temperatures, e.g, the sharp first order phase transition-type change in the APD value about a critical DI, which Fenton et al. (2013) identify as a sigmoidal shape of the APD restitution curve, and associate with a regime where alternans in suppressed.

Dedicated studies by Filippi et al. (2014), into the mechanisms that induce fibrillation during hypothermia. reveal that prolongation of APD is accompanied by comparable reduction in CV at low temperatures, resulting in effective reduction in spatial wavelength and excitability. Thus, fast pacing frequencies, as those observed with spiral waves in the system, can induce wavebreaks or sustained reentry at low temperatures. Our studies are consistent with these findings. We observed that, a certain range of temperatures (10°−20°*C*) provided the optimal substrate for onset of wavebreak in the remodelled human atria. At these temperatures, the wavelength decreased just enough to simultaneously fit multiple scroll waves in the whole atria. The breakup and regeneration of scroll waves occurred mostly around the PV regions. This is in line with the findings of Chen et al. (2003), who reported the occurrence of triggered electrical activity in single pulmonary vein cardiomyocytes isolated from rabbit hearts, and identified them as possibile underlying causes of proarrhythmicity in the atria. However, contrary to Chen et al. (2003), who observed such behaviour only at high temperatures, our observations were at moderate to low temperatures, which made global cooling ineffective even at 10°*C*. Cooling below 5°*C*, however, delivered sufficient destabilization to the scroll wave, thereby pushing it towards an inexcitable boundary of the domain, favoring termination.

To study the effect of temperature in the whole atria, we performed simulation studies with global cooling. Previous studies by Malki and Zlochiver (2018) in simulated 2D human atria explored the possibility to employ spatial gradients of temperature to enforce spiral wave drifting. Such gradients, though effective in directing the movement of a spiral wave through 2D media, are actually extremely difficult to impose in a real non-uniformly curved 3D system, such as the anatomical atria. In addition, there is the issue of intrinsic electrophysiological heterogeneity that Malki and Zlochiver (2018) neglect in their proof-of-principle study, but which has huge influence on electrical wave propagation, as was demonstrated earlier by Tobón et al. (2013) and now, in our present study.

In another landmark study, Yamazaki et al. (2012) demonstrated for the first time, the possibility to control ventricular arrhythmias in rabbit hearts by applying local cooling. Such studies form the next step in the development of thermal-based arrhythmia management strategies. In particular, it would be interesting to identify specific target regions within the atria, which, when cooled, could lead to highest success rate in removing arrhythmias by destabilization through sudden decrease in tissue excitability. Independently, Guill et al. (2012) studied the effects of local epicardial cooling/heating on the complexity of ventricular arrhythmias. They found that cooling induced deceleration of VF which led to increase in the complexity of VF activation patterns, whereas, heating induced acceleration of VF. In our 3D simulations with the anatomically realistic atria, we have not explored the effects of local heating/cooling. As a first approach, we applied low temperature uniformly to all parts of the atrial wall. In the real heart, blood perfusion and equilibriation with the surrounding body temperature prevents this from happening at least immediately after the cooling is applied. Thus, ideally, one should consider the application of the low temperature on the outer or inner surface of the atrial wall and let the system build its own temperature profile. Given that, cooling down to temperatures like 5°*C* and 10°*C* have such contrasting effects on scroll wave dynamics, one can argue that surface cooling at 5°*C* may not be sufficient to terminate the arrhythmia. However, the atrial wall is very thin. Therefore, elimination of activity from several shells of the atrial wall may actually be sufficient to cause removal of the scroll waves from the entire atria. Furthermore, considering the temperature-dependency of the atrial geometry itself, Fedorov et al. (2008) reported a possible variation, with temperature, of the anisotropy ratio of fibers in the hearts of rabbits and ground squirrels. In our studies, we did not factor in the component of a temperature-dependent anisotropy ratio, owing to the lack of availability of experimental data to base our model on. These paths remain to be explored in future studies.

## AUTHOR CONTRIBUTIONS

RM performed 3D simulations and drafted the manuscript. ANMN performed single cell and 1D simulations. YW visualized data and aided 3D simulations. RM, AVP, EB and YW developed the concept, performed data analysis, and reviewed the manuscript.

## ACKNOWLEDGMENTS

Computational resources from HPC at Max Planck Institute for Dynamics and Self-Organization are appreciated.

## SUPPLEMENTAL DATA

### DATA AVAILABILITY STATEMENT

All datasets generated and analyzed for this study are included in the manuscript.

